# Hepatic pyruvate and alanine metabolism are critical and complementary for maintenance of antioxidant capacity and resistance to oxidative insult in mice

**DOI:** 10.1101/2022.06.14.495517

**Authors:** Nicole K.H. Yiew, Joel H. Vazquez, Michael R. Martino, Stefanie Kennon-McGill, Jake R. Price, Felicia D. Allard, Eric U. Yee, Laura P. James, Kyle S. McCommis, Brian N. Finck, Mitchell R. McGill

## Abstract

Pyruvate is a critical intermediary metabolite in gluconeogenesis, lipogenesis, as well as NADH production. As a result, there is growing interest in targeting the mitochondrial pyruvate carrier (MPC) complex in liver and metabolic diseases. However, recent in vitro data indicate that MPC inhibition diverts glutamine/glutamate away from glutathione synthesis and toward glutaminolysis to compensate for loss of pyruvate oxidation, possibly sensitizing cells to oxidative insult. Here, we explored this using the clinically relevant acetaminophen (APAP) overdose model of acute liver injury, which is driven by oxidative stress. We report that MPC inhibition does indeed sensitize the liver to APAP-induced injury in vivo, but only with concomitant loss of alanine aminotransferase 2 (ALT2). Pharmacologic and genetic manipulation of neither MPC2 nor ALT2 alone affected APAP toxicity, but liver-specific double knockout (DKO) of these proteins significantly worsened the liver damage. Further investigation confirmed that DKO impaired glutathione synthesis and increased urea cycle flux, consistent with increased glutaminolysis. Furthermore, APAP toxicity was exacerbated by inhibition of both the MPC and ALT in vitro. Thus, increased glutaminolysis and susceptibility to oxidative stress requires loss of both the MPC and ALT2 in vivo and exacerbates them in vitro. Finally, induction of ALT2 reduced APAP-induced injury.

## INTRODUCTION

Acetaminophen (APAP) is one of the most widely used antipyretic and analgesic drugs in the world. In a cross-sectional survey of the US population, for example, >20% of both adults and children reported use of APAP at least once in the preceding seven days (1, 2) and similar data have been published from other countries (3–5). However, APAP overdose can cause severe liver injury leading to acute liver failure (ALF) and death (6).

Oxidative stress is believed to be the central driver of APAP hepatotoxicity (7–9). Cytochrome P450 enzymes (P450s), particularly CYP2E1, convert APAP to the reactive intermediate *N*-acetyl-*p*-benzoquinone imine (NAPQI) (10, 11) which depletes the antioxidant glutathione and then binds to mitochondrial proteins (7, 12–14). The mitochondrial protein alkylation coincides with reduced respiration (15) and an increase in mitochondrial reactive oxygen species (ROS) production (7, 16). While it has long been assumed that NAPQI binding to electron transport chain (ETC) complexes causes these effects, that has never been mechanistically proven and therefore remains an open question. In any case, the redox imbalance that results from combined glutathione depletion and increased ROS rapidly promotes phosphorylation and activation of the c-Jun N-terminal kinases (JNK) 1/2 (17, 18), which translocate to mitochondria and further reduce mitochondrial respiration via SHP1-mediated inhibition of Src (19, 20). Other kinases are also activated by the oxidative stress and may exacerbate the injury (21–23). Eventually, the mitochondrial membrane permeability transition occurs (24), mitochondria swell, and their membranes rupture (25). The latter releases intermembrane and matrix proteins into the cytosol, including the endonucleases apoptosis-inducing factor and endonuclease G which then cleave nuclear DNA (26). Ultimately, the damaged hepatocytes die by necrosis (27, 28). Thus, anything that upsets mitochondrial metabolism or cellular redox regulation would be expected to alter APAP toxicity.

Pyruvate is a critical cataplerotic and anaplerotic substrate of the mitochondrial TCA cycle with important roles in gluconeogenesis, lipid synthesis, and NADH generation for the electron transport chain (ETC) (29). Most pyruvate is produced in the cytosol as a product of glycolysis or conversion from lactate and is taken up by mitochondria via the mitochondrial pyruvate carrier (MPC) complex. Interestingly, in cultured cells and in hepatic tumors, genetic deletion of the MPC or pharmacological MPC inhibition not only blocks pyruvate oxidation (30, 31), but also decreases glutathione synthesis and disrupts redox balance (32, 33). Thus, due to the role of mitochondrial dysfunction in APAP-induced liver injury and the critical importance of glutathione as a nucleophile and antioxidant in resistance to APAP, we hypothesized that preventing mitochondrial pyruvate uptake by inhibition or liver-specific deletion of the MPC would worsen mitochondrial damage and oxidative stress after APAP overdose and therefore exacerbate APAP toxicity.

## RESULTS

### MPC inhibition and deletion have no effect on APAP hepatotoxicity

To begin testing our hypothesis, we post-treated mice with 30 mg/kg MSDC-0602 (MSDC), a well-established MPC inhibitor (34, 35), 2 hours after APAP overdose. This dose of MSDC has been shown to alter hepatocellular metabolism (36) and the post-treatment regimen was chosen to avoid an effect on NAPQI formation which is necessary for APAP toxicity and peaks around 1-2 hours after APAP treatment (12). We also co-treated primary mouse hepatocytes with 10 mM APAP and either vehicle or 25 μM MSDC-0602, a concentration known to bind to the MPC and markedly inhibit pyruvate flux in cell cultures (34, 35). However, treatment with MSDC-0602 had no effect on serum ALT activity nor on LDH release, both markers of acute liver injury, from hepatocytes following APAP treatment (Fig. 1A,B).

**Figure 1.**
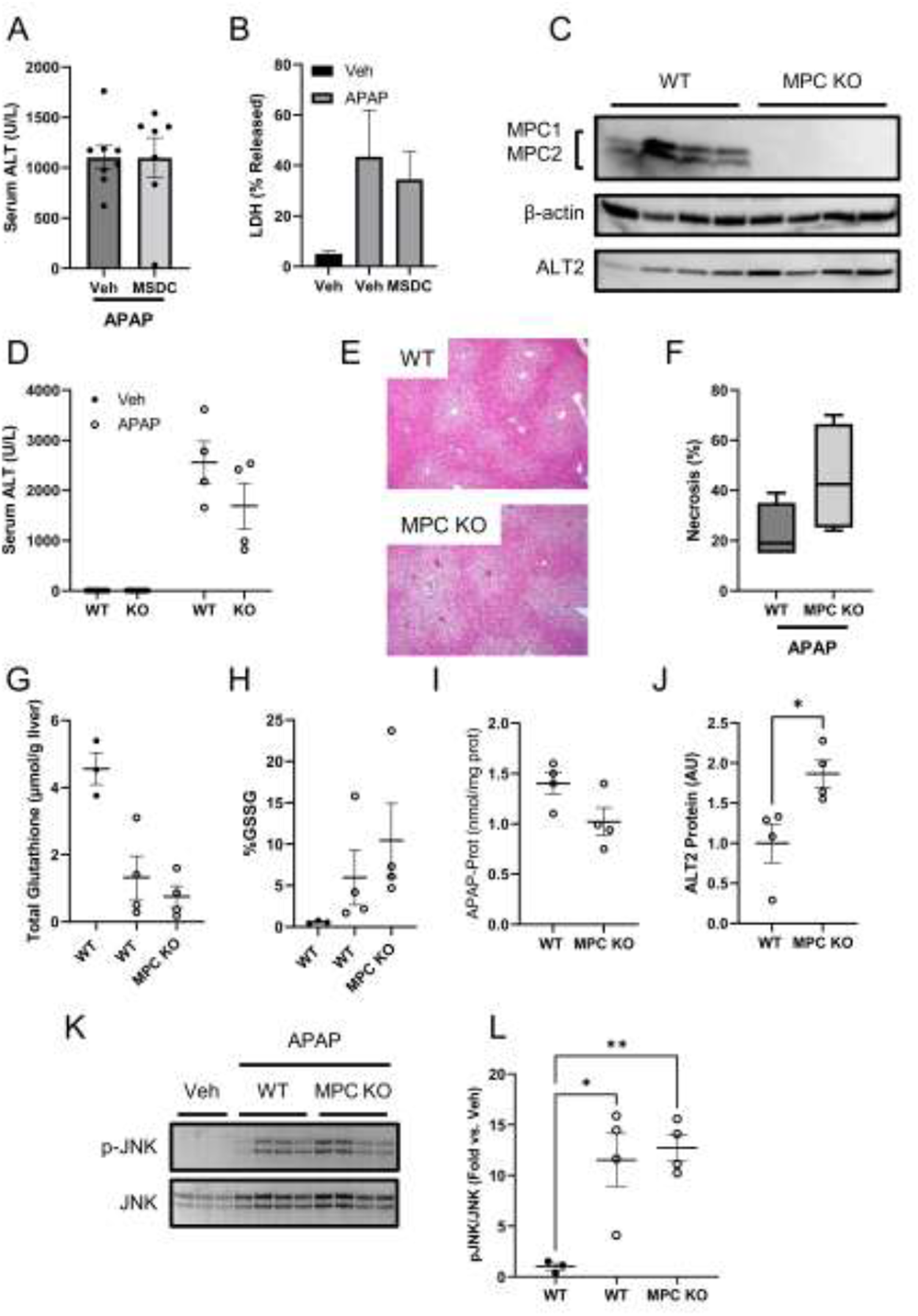
MPC inhibition does not worsen liver injury or glutathione depletion after APAP overdose. (A) Serum ALT from wildtype (WT) mice treated with 30 mg/kg MSDC-0602 (MSDC) or vehicle (Veh) 2 hours after a 300 mg/kg dose of APAP. (B) LDH released from primary mouse hepatocytes co-treated with 10 mM APAP and 25 μM MSDC (n = 3 independent experiments performed in replicates on separate days). (C) Immunoblots for MPC1, MPC2, β-actin, and ALT2 in liver tissue homogenates from liver-specific MPC knockout (KO) mice and littermate wildtype (WT) controls. (D) Serum ALT from MPC KO mice and WT littermates after APAP overdose. (E) H&E-stained liver sections from KO and WT mice after APAP. (F) Box plots of necrosis quantification from liver sections. (G) Total glutathione in liver tissue from KO and WT mice after APAP. (H) Oxidized glutathione in liver tissue from KO and WT mice after APAP. (I) APAP-protein adducts in liver tissue from KO and WT mice (not significant). (J) Densitometric analysis of ALT2 from C. (K) Immunoblots for phosphorylated JNK (p-JNK) and total JNK in liver tissue from KO and WT mice after APAP or Veh. (L) Densitometric analysis of K. Dots show individual values. Lines indicate mean±SE. Box plots show medians and interquartile ranges. *p<0.05, **p<0.01 for the indicated comparisons.

The MPC is composed of two proteins (MPC1 and MPC2) that form a heterodimeric complex in the inner mitochondrial membrane. Because chemical inhibition may not have fully blocked MPC mitochondrial pyruvate flux, we next treated liver-specific MPC2 KO mice with APAP. Consistent with prior reports (35, 37), MPC2 deletion resulted in instability and degradation of MPC1 (Fig. 1C), effectively leading to complete loss of the MPC complex. To our surprise, however, MPC deficiency had no effect on APAP hepatotoxicity at 6 h after treatment, when mitochondrial metabolism is most critical (7, 8). There were no significant differences in serum ALT or hepatic necrosis, total glutathione, oxidized glutathione, APAP-protein binding, or JNK activation between MPC2 KO mice and wildtype littermate controls (Fig. 1D-L).

Although uptake via the MPC is the primary source of mitochondrial pyruvate, it is possible that other metabolic pathways can compensate when the MPC is compromised. In fact, we have previously shown that cytosolic and mitochondrial alanine-pyruvate cycling catalyzed by alanine transaminase enzymes can partially circumvent the loss of MPC activity (35). Indeed, shRNA-mediated knockdown of alanine transaminase 2 (ALT2), which catalyzes the transamination of alanine and α-ketoglutarate to form pyruvate and glutamate in the mitochondrial matrix, further attenuates mitochondrial pyruvate flux in MPC2 deficient hepatocytes (35). In the present study, we found that ALT2 protein abundance was 1.9±0.2-fold greater in livers from MPC2 KO mice compared to WT mice after APAP treatment (Fig. 1C,J), suggesting that mitochondrial alanine-pyruvate cycling might be compensating for loss of the MPC during APAP hepatotoxicity as well.

### ALT2 inhibition and deletion have no effect on APAP hepatotoxicity

To test the hypothesis that ALT2 has a role in APAP-induced liver injury, we first used a pharmacological approach. β-chloro-*l*-alanine (BCLA) is an alanine analog and suicide substrate of ALT that irreversibly inhibits the enzyme’s activity (38). Although it has been extensively used in studies with bacteria and yeast, there are no data in the literature to guide its application in mice. Thus, we first performed timing and dose-response studies. The results revealed that BCLA can inhibit ALT activity in the liver *in vivo*, the effect is greater at 24 h post-BCLA treatment than an earlier time point, and ≥50% inhibition can be achieved with a dose of 100 mg/kg (Suppl. Fig. 1). Based on these data, we adopted a 24 h single-dose (Fig. 2A-D) and 48 h two dose (Fig. 2E,F) pre-treatment regimen with 100 mg/kg BCLA to study the role of ALT in APAP toxicity. BCLA reduced both serum and hepatic ALT activity after APAP overdose (Fig. 2A,B,E). However, like MSDC and MPC2 deletion, BCLA had no effect on injury as indicated by serum LDH and hepatic necrosis (Fig. 2C-D).

**Figure 2.**
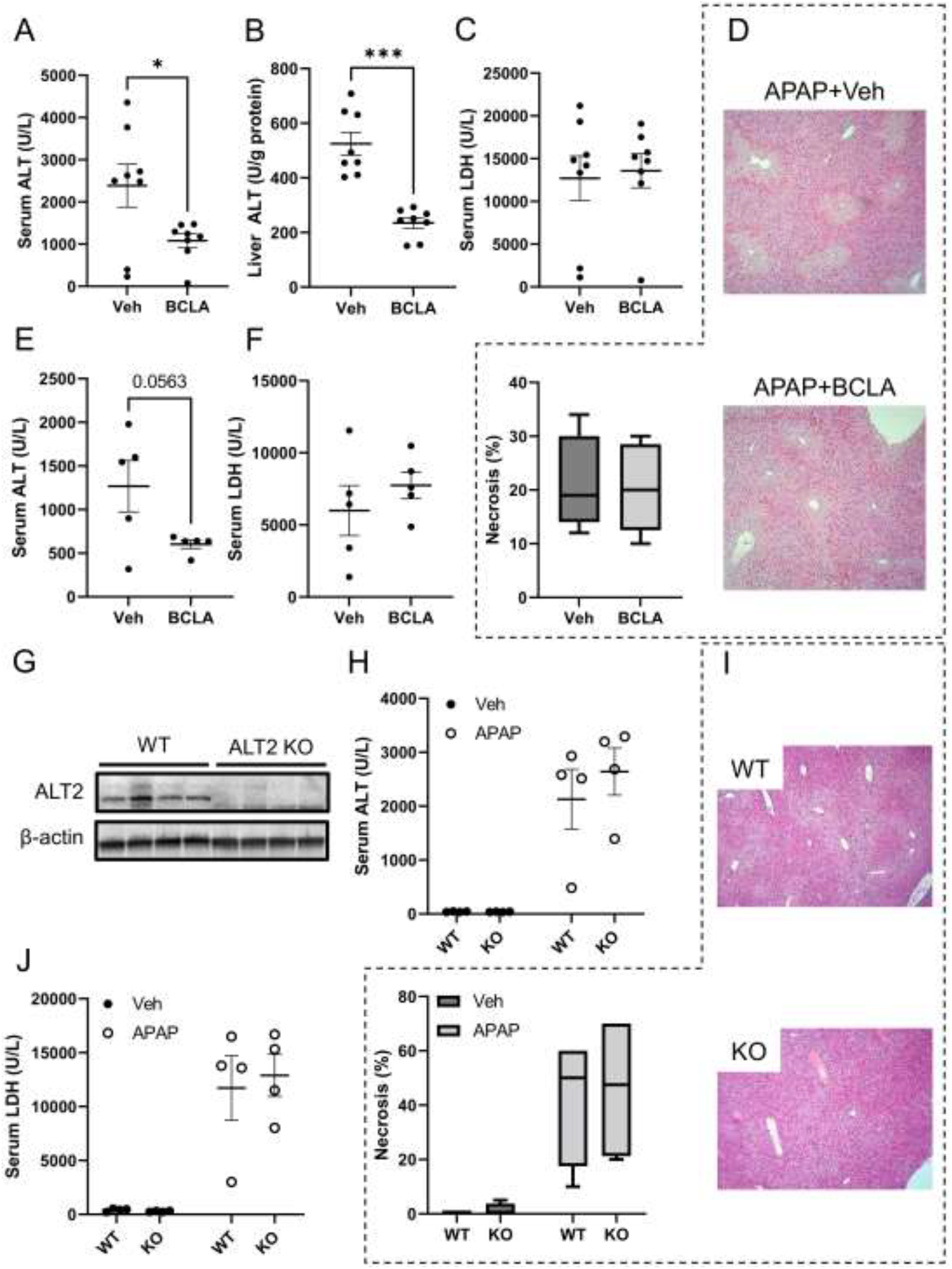
ALT2 inhibition does not worsen liver injury after APAP overdose. (A-D) WT mice were pre-treated with 100 mg/kg β-chloro-*l*-alanine (BCLA) or vehicle (Veh) then received a 300 mg/kg dose of APAP 24 h later. (E-F) WT mice were pre-treated with 100 mg/kg β-chloro-*l*-alanine (BCLA) or vehicle (Veh) once per day for 2 days then received a 300 mg/kg dose of APAP on the third day. (G-J) Liver-specific ALT2 knockout (KO) mice and wildtype (WT) littermate controls were treated with 300 mg/kg APAP. (A,E,H) Serum ALT. (B) Liver ALT. (C,F,J) Serum LDH. (D,I) H&E-stained liver sections with necrosis quantification. (G) Immunoblots for ALT2 and β-actin in liver tissue from WT and KO mice. Dots show individual values. Lines indicate mean±SE. Box plots show medians and interquartile ranges. *p<0.05, ***p<0.001 for the indicated comparisons.

BCLA did not completely block hepatic ALT activity (Fig. 2B). In addition, it is possible that the drug cannot reach ALT2 within the mitochondrial matrix. Thus, to ensure complete inhibition of ALT2, we next used liver-specific ALT2 KO mice (Fig. 2G). However, like MPC2 KO, there was significant effect of ALT2 deletion on APAP-induced liver injury, as indicated by serum ALT (Fig. 2H) and LDH (Fig. 2J) and by hepatic necrosis (Fig. 2I). Although it may seem surprising that ALT2 KO itself did not reduce serum ALT activity, this observation is consistent with reports that ALT1 is the dominant isoform released into serum (39–41), accounting for ≥90% after injury (41). Thus, we continued to use serum ALT as our primary measure of liver injury. Together, these data indicate that neither MPC2 deletion nor ALT2 deficiency alone alters APAP hepatotoxicity but do not rule out the possibility of complementary effects.

### Combined MPC2/ALT2 deficiency worsens APAP-induced liver injury

To determine if deletion of both MPC2 and ALT2 worsened APAP hepatotoxicity, we generated liver-specific ALT2/MPC2 double KO (DKO) mice (Suppl. Fig. 2). Consistent with our hypothesis that loss of both proteins more significantly reduced mitochondrial pyruvate availability and altered mitochondrial metabolism in APAP toxicity, DKO mice had greater serum ALT activity and hepatic necrosis after APAP overdose (Fig. 3A-C). Importantly, they also had lower total hepatic glutathione levels (Fig. 3D). Glutathione reaches its nadir with nearly complete loss by approximately 0.5 to 1 hour post-APAP and then recovers over the following 5-6 hours (12), so these 6 hour data suggest impaired glutathione re-synthesis, likely due to the anaplerotic diversion of glutamate to α-ketoglutarate (αKG) and away from glutathione synthesis. Furthermore, these effects occurred despite less APAP-protein binding in the DKO mice (Fig. 3E). Because protein alkylation is the initiating event in APAP hepatotoxicity, even greater injury may have been observed had the level of protein binding been the same as in WT. Indeed, when we normalized serum ALT to hepatic APAP-protein adducts, the difference in means was even greater with a 4.9±0.2-fold increase in ALT in the DKO animals (mean±SE) (Fig. 3F). The reason for the reduced adducts in DKO animals was likely depletion of NADPH, which is necessary for the P450 catalysis that produces NAPQI. The protein levels of CYP2E1 – the primary enzyme that converts APAP to NAPQI (42) – were the same between genotypes (Fig. G,H), demonstrating that the lower NAPQI production and protein binding was not due to altered CYP2E1 expression. Furthermore, lower NADPH levels have been reported in MPC1-deficient cells *in vitro* (33) and we observed NADPH depletion in our DKO animals after APAP overdose as well (Fig. 3I,J). Finally, the DKO animals experienced greater oxidative stress (Fig. 3K). To determine if the opposite approach – increasing mitochondrial pyruvate flux rather than completely blocking it – might protect against the injury, we pre-treated WT mice for 24 h with a dose of dexamethasone known to induce ALT2 expression (43, 44) followed by APAP. Western blotting confirmed an increase in hepatic ALT2 and histology revealed that it was indeed protective (Fig. 3L-N).

**Figure 3.**
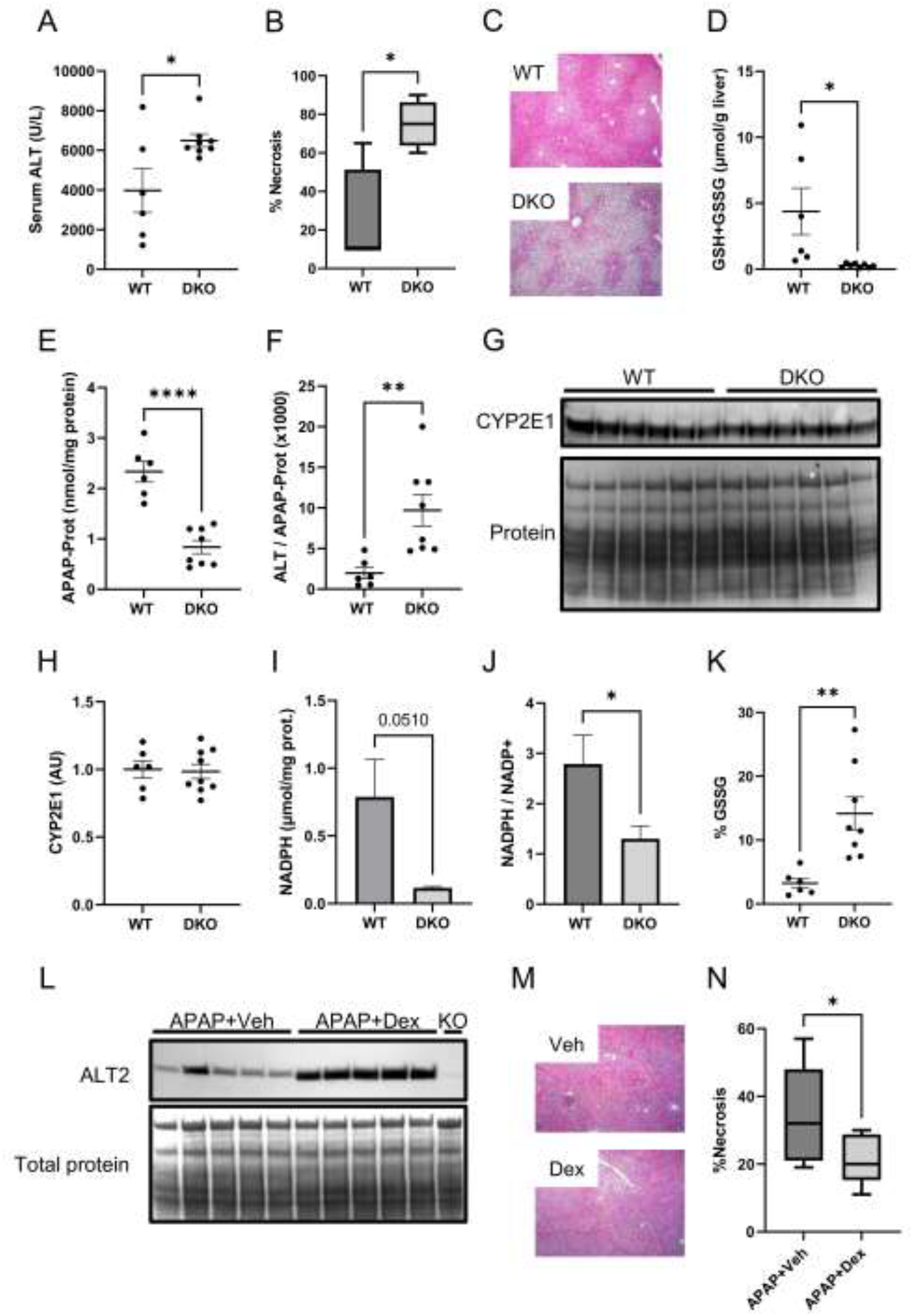
MPC2/ALT2 double-knockout signficantly worsens liver injury and prevents glutathione recovery after APAP overdose. Mice with liver-specific double knockout (DKO) of MPC2 and ALT2 and wildtype (WT) littermate controls were treated with 300 mg/kg APAP. (A) Serum ALT. (B) Necrosis quantification. (C) H&E-stained liver sections. (D) Hepatic total (GSH+GSSG) glutathione. (E) Hepatic APAP-protein adducts. (F) Ratio of serum ALT to APAP-protein adducts. (G) Immunoblot for CYP2E1 and total protein from liver tissue. (H) Densitometric analysis of G. (I) Hepatic NADPH concentration. (J) Ratio of NADPH to NADP^+^ in the liver. (K) Percentage of hepatic glutathione in the oxidized form (GSSG). Densitometric analysis of K. Dots show individual values. Lines indicate mean±SE. Box plots show medians and interquartile ranges. *p<0.05, **p<0.01, ****p<0.0001 for the indicated comparisons.

### The mechanism of exacerbated injury in DKO mice is diversion of glutamate away from glutathione

The reason for impaired glutathione re-synthesis in the DKO mice may be diversion of glutamine/glutamate away from glutathione synthesis and into the TCA cycle to maintain TCA cycle flux. This was previously reported in MPC1-deficient hepatocytes *in vitro* (32) and in other conditions in which mitochondrial pyruvate metabolism was impaired (45). In addition, glutaminolysis and glutamate utilization has been proposed as a source of TCA intermediates in cancer cells (46, 47). Thus, to further test the idea that glutamate is a major anaplerotic substrate in the liver *in vivo* when pyruvate flux is compromised, we compared basal total hepatic glutathione between DKO and WT mice and between HepG2 cells treated with the combination of another MPC inhibitor, UK-5099, and BCLA or vehicle. The concentration of these inhibitors *in vitro* and the treatment regimen were chosen to mimic combined MPC and ALT2 KO.

Interestingly, glutathione was 22±5% (mean±SE) lower in liver tissue from DKO animals (Fig. 4A) and almost undetectable in the UK/BCLA co-treated cells (Fig. 4B). In addition, post-treatment of cells with UK/BCLA after glutathione depletion with buthionine sulfoximine (BSO) completely prevented glutathione re-synthesis over the next 24 h (Fig. 4C). Furthermore, APAP toxicity was dramatically increased in HepG2 cells overexpressing CYP2E1 when APAP was co-administered with UK and BCLA (Fig. 4D-G), similar to the DKO mice. Importantly, CYP2E1-overexpressing HepG2 cells were previously validated for the study of APAP hepatotoxicity, with formation of APAP-protein adducts, mitochondrial damage, and cell death occurring similar to primary mouse and human hepatocytes (48), and our preliminary dose-response and time course data were consistent with those earlier results (Suppl. Fig. 4). Although one might think fatty acid oxidation (FAO) can serve as a source of acetyl-CoA for the TCA cycle and potentially also compensate for MPC loss, FAO is inhibited in APAP toxicity (49–51) and there was no difference in serum ketones between DKO and WT mice in the present study (Suppl. Fig. 5). Furthermore, FAO adds only two carbons to the TCA cycle compared to glutamate’s four and each turn results in loss of two carbons as CO_2_, so the net contribution of FAO is nil, and furthermore increased FAO would be cataplerotic – stealing carbons from the TCA cycle for more fatty acid synthesis. Thus, it is unlikely that FAO compensates during APAP-induced injury. Finally, targeted plasma metabolomics revealed elevated urea cycle intermediates (citrulline, arginine, and ornithine) in untreated DKO mice (Fig. 5), consistent with increased disposal of ammonia from glutamate and other amino acids. Importantly, conversion of oxaloacetate to aspartate before urea cycle entry also consumes glutamate. Thus, increased urea cycle flux may have also contributed to the impaired glutathione synthesis in these animals. Together, these data are consistent with increased conversion of glutamate and other amino acids to α-keto acids to feed the TCA cycle.

**Figure 4.**
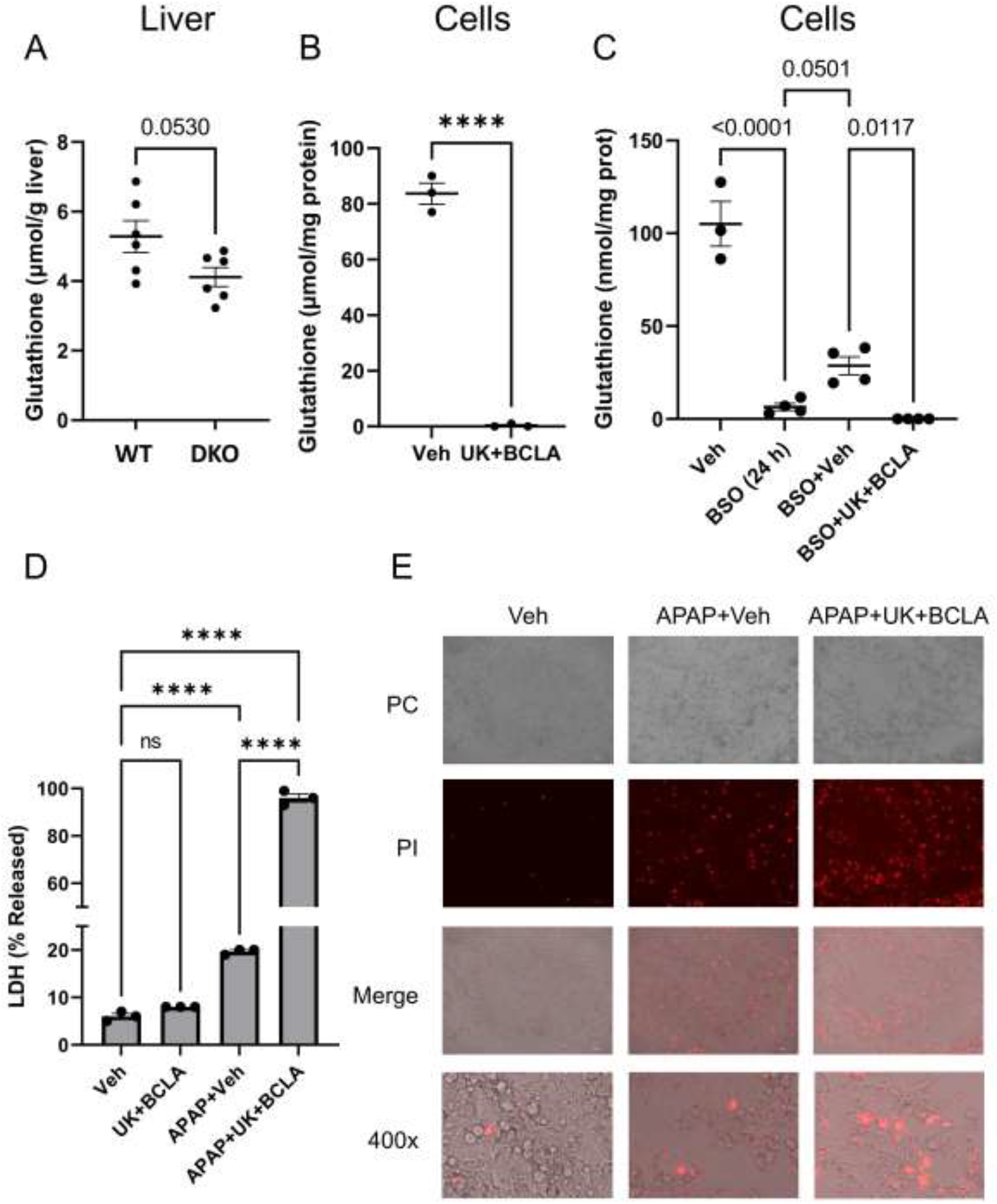
Combined inhibition of the MPC and ALT2 reduces glutathione synthesis *in vivo* and *in vitro*. (A) Hepatic total glutatione in mice with liver-specific double knockout (DKO) of MPC2 and ALT2 and wildtype (WT) littermate controls without treatment. (B) Total glutathione from HepG2 cells co-treated with 25 µM UK-5099 (UK) and 250 µM β-chloro-*l*-alanine (BCLA) or vehicle (Veh) for 24 h. (C) Total glutathione from HepG2 cells treated with Veh or buthionine sulfoximine (BSO) for 24 h followed by 24 h recovery without BSO but with either Veh (BSO+Veh) or UK and BCLA (BSO+UK+BCLA). (D) LDH released from HepG2 cells overexpressing CYP2E1 and treated with Veh, UK+BCLA, 20 mM APAP+Veh, or APAP+UK+BCLA for 48 h. (E) Phase contrast (PC) imaging and propidium iodide (PI) staining cells from D. Dots show individual values. Lines indicate mean±SE. ****p<0.0001 for the indicated comparisons.

**Figure 5.**
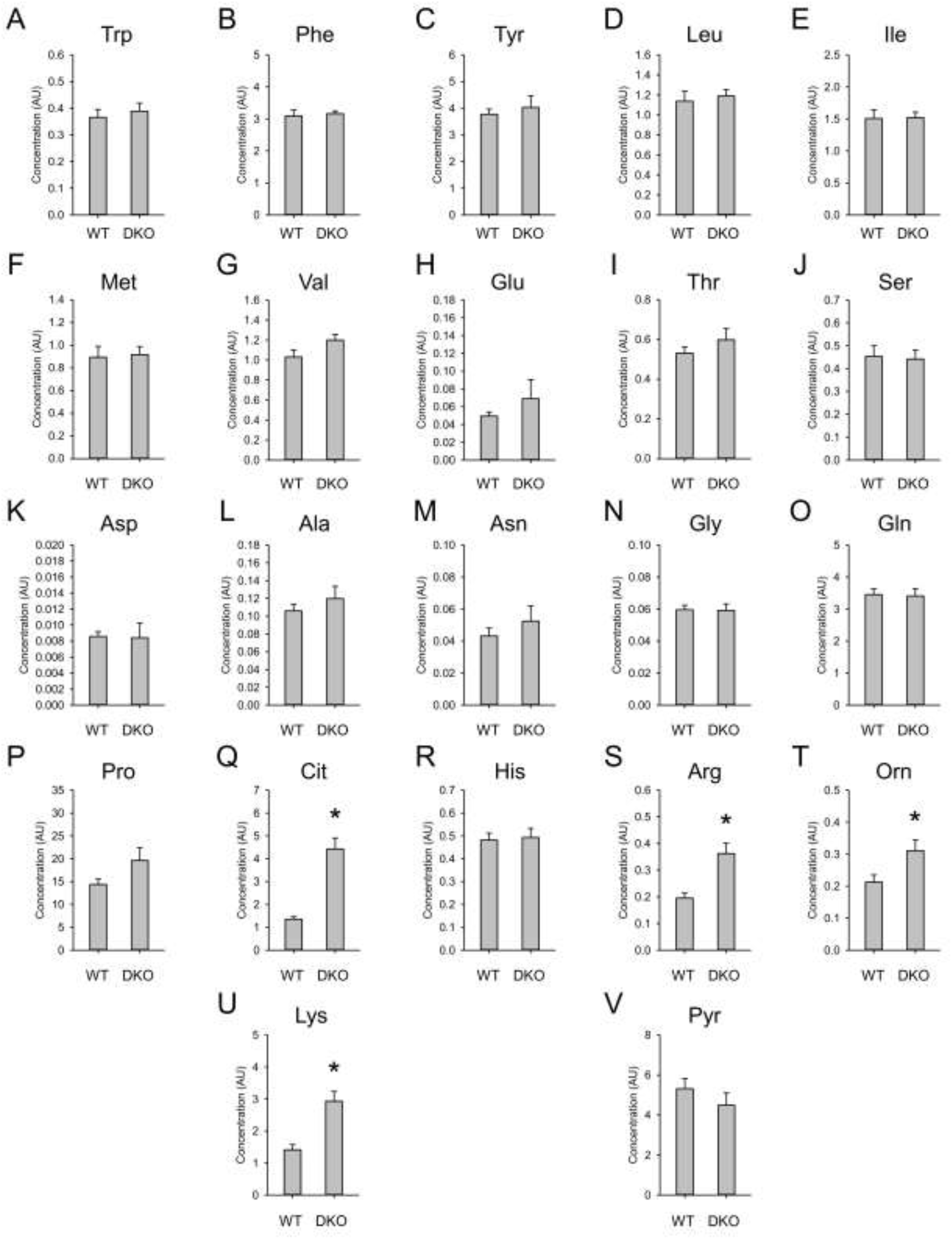
MPC2/ALT2 double-knockout increases urea cycle flux. Amino acids and pyruvate (Pyr) were measured in plasma from mice with liver-specific double knockout (DKO) of MPC2 and ALT2 and wildtype (WT) littermate controls. Graphs show mean±SE. *p<0.05 vs. WT.

### NAC still protects in MPC2 KO and DKO mice

NAC is currently the standard-of-care treatment for APAP overdose in humans. It is generally believed to protect against APAP hepatotoxicity by providing cysteine for re-synthesis of glutathione which can then scavenge NAPQI and ROS. However, the doses of NAC given to APAP overdose patients (140 mg/kg loading, followed by additional lower doses) greatly exceed those required to replenish glutathione. In addition, it has been reported that similar doses can increase flux through the mitochondrial pyruvate dehydrogenase complex (PDHC) in mice (52). It has therefore been proposed that large doses of NAC like those administered clinically protect against APAP in part by increasing pyruvate oxidation (52, 53) – an effect that would require the MPC. To test this hypothesis more directly than in prior work, we examined NAC protection in DKO mice and WT littermate controls. Interestingly, NAC protected equally well in the two genotypes (Fig. 5A), despite using the same large dose (318 mg/kg) employed in the prior work (53). To our surprise, NAC also stimulated glutathione re-synthesis in the DKO mice similar to the WT animals (Fig. 5B). These data suggest that while excess NAC may increase PDHC flux, this is not required for NAC-mediated protection after APAP hepatotoxicity. Rather, some of the NAC-derived cysteine may be converted to glutamate, which can then either enter the TCA cycle and increase TCA cycle flux overall or can be used to re-synthesize glutathione.

## DISCUSSION

Prior research has suggested that APAP overdose leads to impairments in mitochondrial metabolism (15) and that maintaining flux through these pathways can attenuate liver injury in response to this stimulus (53). Evidence has emerged that use of amino acids, particularly alanine and glutamine/glutamate, in the TCA can compensate for MPC deficiency (35, 54, 55). Indeed, it was first reported nearly a decade ago that glutaminolysis (the conversion of glutamine to glutamate to α-KG) constitutes an important anaplerotic input for the TCA cycle when mitochondrial pyruvate metabolism is impeded (54). However, there is concern that increased flux of glutamate to the TCA cycle could lead to decreased glutamate availability for glutathione synthesis. Indeed, more recent studies have illustrated that MPC inhibition leads to impaired glutathione synthesis and regeneration of oxidized glutathione in cultured cells. The first study to show this (32) demonstrated that inhibition of the MPC with UK-5099 moderately decreased glutathione concentrations in Hepa1-6 cells and that MPC1 KO in primary hepatocytes partially impaired glutathione re-synthesis. The second study demonstrated that MPC1 deletion also dramatically increased the GSSG/GSH ratio in C2C12 myocytes, likely by depleting the NADPH required for reduction of GSSG back to GSH via glutathione reductase (33). However, these effects were not observed in liver tissue from MPC1 KO mice *in vivo* (32). Based on our data, the explanation for this discrepancy between cultured cells and mice is likely compensation for loss of MPC activity by ALT2 in the animals. ALT2 protein was roughly 2-fold elevated in our MPC2 KO mice and there was no evidence of worsened glutathione depletion in these animals after APAP overdose. However, we found that when both MPC2 and ALT2 were absent, hepatic glutathione was decreased at baseline compared to WT mice and was dramatically lower after APAP. Furthermore, the phenotype we observed in cells treated with both UK-5099 and the ALT inhibitor BCLA was much more severe than in prior studies using an MPC inhibitor alone. We found complete elimination of basal glutathione and total inhibition of re-synthesis after pharmacologic depletion. Altogether, our results suggest that loss of both the MPC and ALT2 is required to alter glutathione synthesis in the liver *in vivo*. Interestingly, we also found that inducing expression of ALT2 can reduce APAP-induced liver injury, which could be explored in the future as a novel therapeutic approach in liver injury and disease.

There is currently much interest in the development of drugs to target the MPC to treat metabolic diseases. Because pyruvate can be converted to glucose through a series of steps that begins with conversion to oxaloacetate within mitochondria, inhibiting mitochondrial pyruvate uptake could theoretically reduce hyperglycemia and its downstream adverse effects. Indeed, obesity- and streptozotocin-induced hyperglycemia is attenuated in mice with liver-specific MPC KO or with MPC inhibitor treatment (35, 56). MPC inhibitors may also have promise in treating nonalcoholic steatohepatitis, as MPC deletion or inhibition protects mice from liver injury and fibrosis in preclinical models (57, 58). However, given the central importance of pyruvate in intermediary metabolism, there are valid concerns regarding the safety of MPC inhibition in all contexts and tissue types. For example, it is possible that the alterations in glutathione synthesis and redox status observed in cell culture studies with MPC inhibition could predispose patients to oxidative and mitochondrial insults. However, our observation that ALT2 induction can compensate for MPC loss *in vivo* – avoiding increased glutaminolysis and sparing glutathione – argues against that idea in the liver.

Although our data demonstrate that diversion of glutamine/glutamate to the TCA cycle compromises glutathione re-synthesis after APAP specifically in the extreme case of MPC and ALT2 DKO, there is some evidence that this could occur to a lesser extent even in WT animals as well. It was recently demonstrated that mice deficient in pyruvate dehydrogenase kinase 4 (PDK4), which phosphorylates and inhibits the PDHC, had much lower liver injury, oxidative stress, and mitochondrial JNK translocation after APAP overdose than WT controls (59). They also observed greater glutathione re-synthesis in the PDK4-deficient mice. This effect on glutathione could have been due to lower injury and therefore faster recovery, but it could also have been due in part to the glutathione-sparing effect of increased pyruvate oxidation. If so, then our data and theirs suggest a continuum, with our DKO mice and their PDK4 KO mice at opposite extremes and the WT condition somewhere in the middle. That is supported by the fact that dexamethasone pre-treatment induced ALT2 and reduced APAP-induced liver injury in our WT animals.

NAC is the current standard-of-care treatment for APAP overdose. The primary mechanism by which NAC protects against APAP hepatotoxicity is providing cysteine for the rapid re-synthesis of glutathione (14, 60, 61). However, it is often claimed that NAC is beneficial in APAP overdose patients even when administered late after overdose, when APAP bioactivation and ROS production have already occurred (62, 63). There is also modest evidence that NAC is beneficial for patients with non-APAP acute liver damage presumably unaccompanied by glutathione depletion (64). One of the most compelling hypotheses to explain how NAC protects in these scenarios is increased pyruvate oxidation. Zwingmann and Bilodeau demonstrated that NAC increases glucose flux through the PDHC and TCA cycle and protects against oxidative stress even in glutathione-replete mice (52). The mechanism by which NAC brought about this effect was not clear from that study but could have been conversion of the excess cysteine to pyruvate. It was later demonstrated that when equimolar doses of NAC and the tripeptide glutathione were administered to mice after APAP overdose, glutathione was more effective than NAC at reducing APAP-induced liver injury and it increased TCA cycle flux to a greater degree (53). Furthermore, increasing the NAC dose 3-fold to provide the same number of amino acids as glutathione equalized their effects (53). Based on this, it was concluded instead that amino acids derived from NAC may be used to synthesize TCA cycle substrates which then enhance TCA flux. This hypothesis is supported by our observation that the same large dose of NAC used in those studies protected equally well in WT and DKO mice that cannot use cytosolic cysteine-derived pyruvate. In addition, the fact that NAC also restored hepatic glutathione in DKO mice is consistent with the possibility that some of the excess cysteine was instead converted to glutamate, which can then be used in both glutathione re-synthesis and the TCA cycle.

Finally, ALT activity has been the primary serum biomarker of liver injury since 1955 (65, 66). However, the enzyme also catalyzes an important reaction in intermediary metabolism. Despite this, its metabolic and physiological roles in liver injury and disease are rarely considered. Interestingly, it was recently reported that hepatic ALT1 and ALT2 contribute to hyperglycemia in diabetic mice by converting alanine to pyruvate which is then used in gluconeogenesis (67, 68), providing possibly the first evidence that ALT2 has pathophysiological significance in liver disease. The current study adds to our emerging awareness that ALT is more than a passive biomarker of injury by demonstrating its complementary role with the MPC in maintenance of mitochondrial intermediary metabolism and preservation of glutathione levels.

## CONCLUSIONS

Our data suggest that hepatocytes have metabolic redundancies to preserve the TCA cycle flux and defend against oxidative stress even in the context of impaired mitochondrial pyruvate metabolism (Fig. 6), which is an important cataplerotic and anaplerotic metabolic substrate of the TCA cycle. We found that when mitochondrial pyruvate import is compromised, ALT2-mediated pyruvate synthesis in the mitochondrial matrix can largely compensate. However, if all sources of mitochondrial pyruvate are impeded, hepatocytes likely switch to metabolism of other amino acids to maintain flux, but at the cost of reduced synthesis of the antioxidant glutathione. As the amino acid glutamate is shunted into the TCA and urea cycles, less is available for glutathione synthesis, rendering hepatocytes more susceptible to oxidative insult. In addition, we also found that NAC still protects against APAP hepatotoxicity in DKO mice. These findings have implications for the safety and future development of MPC inhibitors to treat metabolic diseases and for our understanding of the mechanisms by which NAC protects from liver injury following APAP overdose.

**Figure 6.**
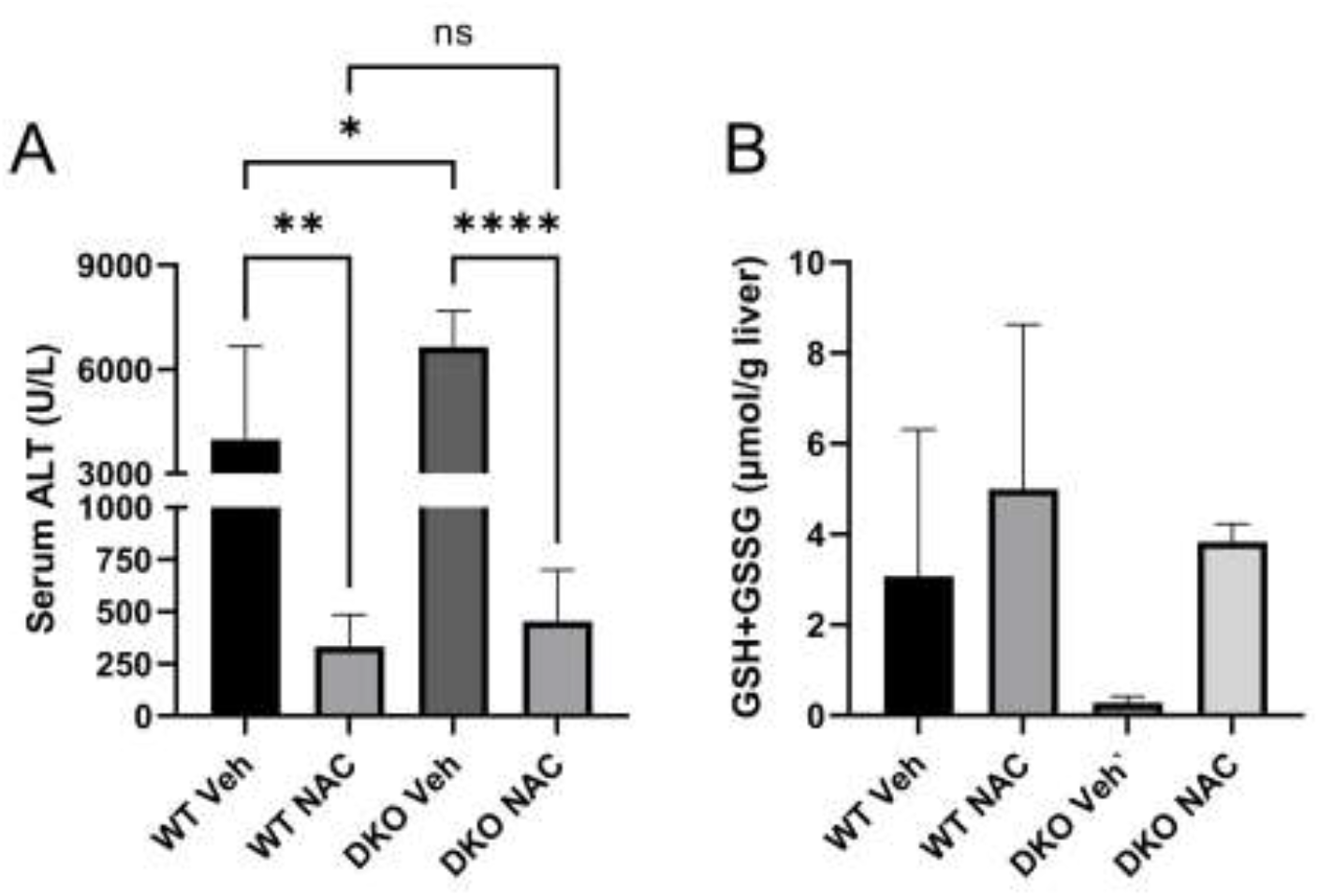
N-acetyl-l-cysteine still protects in MPC2/ALT2 double-knockout mice. Mice with liver-specific double knockout (DKO) of MPC2 and ALT2 and wildtype (WT) littermate controls were treated with vehicle (Veh) or 300 mg/kg APAP intraperitoneally followed by 318 mg/kg *N*-acetyl-*l*-cysteine or Veh intravenously 2 h later. (A) Serum ALT. (B) Hepatic total glutathione (GSH+GSSG). Graphs show mean±SE. *p<0.05, **p<0.01, ****p<0.0001 for the indicated comparisons.

**Figure 7.**
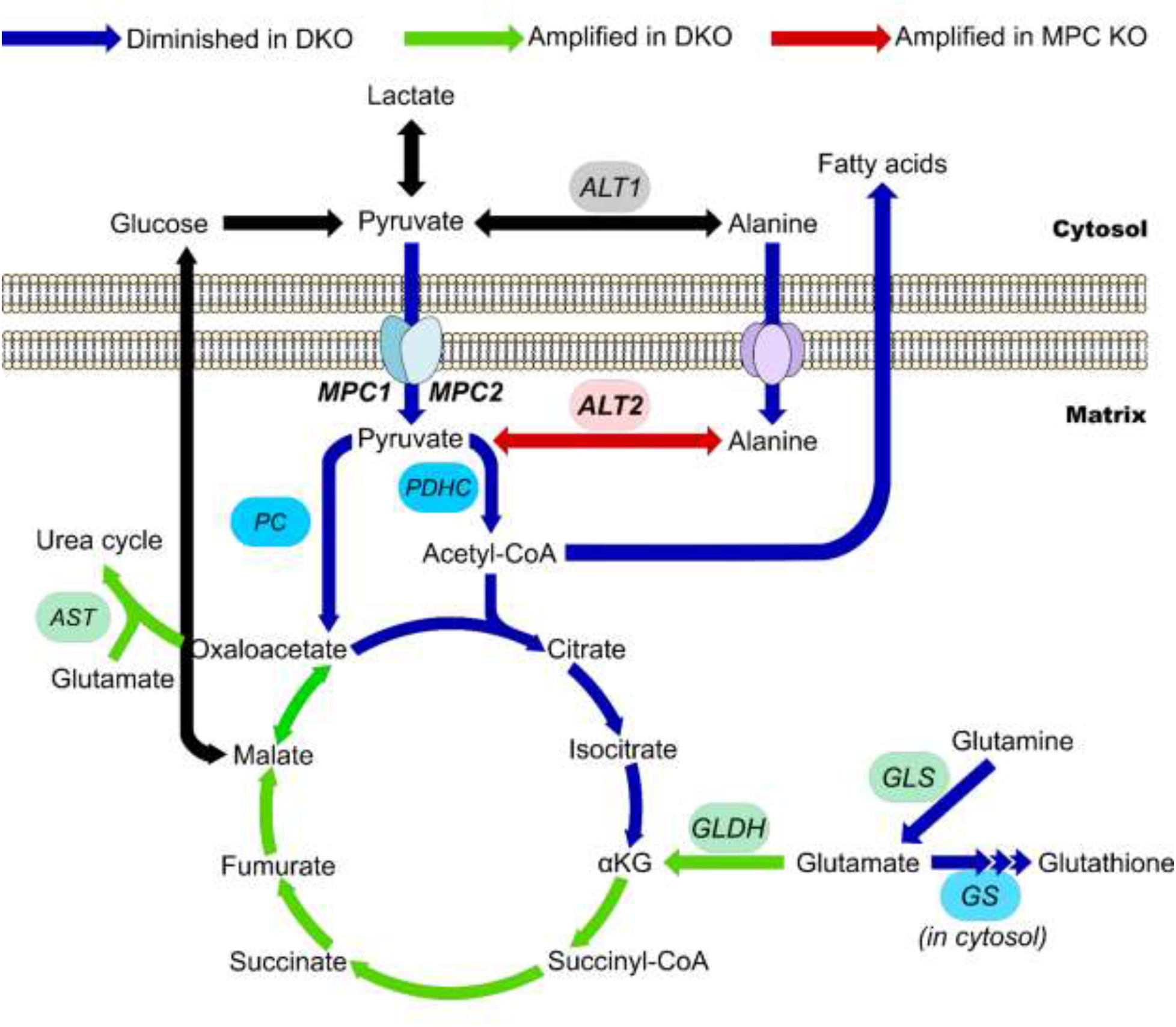
Schematic representing tricarboxylic acid cycle flux early during APAP toxicity in the different genotypes tested. In wildtype (WT) mice, the tricarboxylic acid (TCA) cycle and mitochondrial metabolism more generally proceeds through all arrows. Blue arrows indicate pathways or steps that are diminished in mice with liver-specific double knockout (DKO) of MPC2 and ALT2. Green arrows indicate pathways that are amplified in the DKO animals. Red arrows indicate pathways or steps that are amplified specifically in MPC2 KO mice. αKG, α-ketoglutarate. ALT, alanine aminotransferase. AST, aspartate aminotransferase. GLDH, glutamate dehydrogenase. GLS, glutaminase. GS, glutathione synthase. MPC, mitochondrial pyruvate carrier. PC, pyruvate carboxylase. PDHC, pyruvate dehydrogenase complex.

## EXPERIMENTAL PROCEDURES

### Animals

WT male C57BL/6J mice between the ages of 7 and 8 weeks were purchased from the Jackson Laboratory (Bar Harbor, ME, USA).

The generation of mice harboring a floxed allele of the gene encoding ALT2 (*Gpt2*) derived from *Gpt2* germline KO mice generated by the Knockout Mouse Project Repository (KOMP) (project ID CSD24977) has recently been described (68). Similarly, the generation of *Mpc2* floxed mice using ES cells obtained from the European Conditional Mouse Mutagenesis Program (EUCOMM) (International Knockout Mouse Consortium [IKMC] project 89918, catalog no. MAE-1900) has also been described (35). Transgenic mice expressing *Cre* under control of the albumin promoter (B6.Cg-Speer6-ps1Tg(Alb-cre)21Mgn/J; Jackson Laboratory stock number: 003574) were bred with these animals to create liver-specific KO mice. Combined liver-specific ALT2/MPC2 DKO mice were then generated by intercrossing the two liver-specific single gene knockouts. Littermate mice not expressing *Cre* (fl/fl mice) were used as controls for all liver knockout studies.

Animals were kept in a temperature-controlled 12 h light/dark cycle room with *ad libitum* access to food and water. For BCLA time course and dose-response experiments, fed WT mice were treated with BCLA dissolved in 1x phosphate-buffered saline (PBS) at the indicated doses or PBS vehicle alone and liver tissue was collected at the indicated time points. For APAP experiments, mice were fasted overnight, then treated i.p. with 300 mg APAP per kg body mass (mg/kg) dissolved in 1x phosphate-buffered saline (PBS). Some WT mice were pre-treated i.p. with the specified doses of BCLA or PBS vehicle for the indicated times before APAP. Other received 30 mg/kg MSDC, 318 mg/kg NAC, or appropriate vehicle 2 h after APAP. NAC and NAC vehicle (0.9% saline) were administered i.v. All other treatments were given i.p. Blood and liver tissue were collected at 6 h post-APAP. Some liver tissue pieces were fixed in 10% phosphate-buffered formalin for histology, while others were flash-frozen in liquid nitrogen and stored at −80°C for later biochemical analyses. All study protocols were reviewed and approved by the Institutional Animal Care and Use Committees of the University of Arkansas for Medical Sciences and Washington University in St. Louis and are consistent with best practices in the *Guide for the Care and Use of Laboratory Animals*.

### Biochemistry

Alanine aminotransferase (ALT) was measured in serum using a kit from MedTest Dx, Inc. (Canton, MI, USA), according to the manufacturer’s instructions. Lactate dehydrogenase (LDH) was measured in serum as previously described (69). For tissue ALT activity, frozen liver tissue was homogenized in 25 mM HEPES buffer with 5 mM EDTA (pH 7.4), 0.1% CHAPS, and protease inhibitors (luepeptin, pepstatin A, and aprotinin) using a bead homogenizer (Thermo Fisher Scientific, Waltham, MA). After centrifugation at 10,000 x *g* and 4°C for 5 min, ALT activity was measured in the supernatant using the kit from Pointe Scientific, Inc., mentioned above. Protein concentration in the lysates was measured using the bicinchoninic acid (BCA) assay and the values were used to normalize hepatic ALT activity.

### Histology

Formalin-fixed tissue was embedded in paraffin wax and 5 µm sections were mounted on glass slides for hematoxylin and eosin (H&E) staining and immunohistochemistry. H&E staining was performed using a standard protocol.

### Western blotting

Western blotting was performed as we previously described (70). Briefly, liver tissue was homogenized in 25 mmol/L HEPES buffer with 5 mmol/L EDTA, 0.1% CHAPS, and a mixture of the protease inhibitors leupeptin, pepstatin A, and aprotinin (pH 7.4; Sigma). Protein concentration was then measured using the BCA assay. After dilution in homogenization buffer, the samples were mixed with reduced Laemmli buffer (Bioworld, Dublin, OH, USA) and boiled for 1 min. Equal amounts of protein (50 μg) were added to each lane of a 4%–20% Tris-glycine gradient gel. After electrophoresis separation, proteins were transferred to polyvinylidene fluoride (PVDF) membranes and blocked with 5% milk in Tris-buffered saline with 0.1% Tween 20 (TBS-T). The primary antibodies against phospho-JNK (Cat. No. 4668), JNK (Cat. No. 9252), and MPC2 (Cat. No. 46141) were purchased from Cell Signaling Technology (Danvers, MA, USA) and used at 1:1,000 dilution in TBS-T with 5% milk. The Cyp2e1 antibody was purchased from Proteintech (Rosemont, IL, USA) (Cat. No. 19937-1-AP). The ALT2 antibody was from Sigma (St. Louis, MO, USA) (Cat. No. HPA051514). Secondary antibodies were purchased from Li-Cor Biotechnology (Lincoln, NE, USA) and Cell Signaling Technology. Total protein was stained with Coomassie blue. All blots were imaged using a Li-Cor Odyssey instrument (phospho-JNK and JNK) or an Invitrogen iBright FL1500 (all other immunoblots and Coomassie stained membranes).

### Plasma metabolomics for amino acids and pyruvate

Amino acids were extracted from 20 μL of plasma with 200 μL of methanol containing Try-d8 (1600ng), Phe-d8 (160ng), Tyr-d4 (160ng), Leu-d3 (320ng), Ile-13C6,15N (160ng), Met-d3 (160ng), Val-d8 (400ng), Glu-d3 (2400ng), Thr-13C4 (320ng), Ser-d3 (400ng), Asp-d3 (3200ng), Ala-d4 (1600ng), Asn d3,15N2(3200ng), Gly-d2 (16000ng), Gln-13C5 (3200ng), Pro-d7 (800ng), Cit-d4 (80ng), His 13C6 (400ng), Arg-13C6 (1600ng), Orn-13C5 (400ng), Lys-d8 (400ng), and pyruvic acid-13C5 (80 ng) as the internal standards for Try, Phe, Tyr, Leu, Ile, Met, Val, Glu, Thr, Ser, Asp, Ala, Asn, Gly, Gln, Pro, Cit, His, Arg, Orn, Lys, and pyruvic acid (Pyr) respectively. The sample aliquots for pyruvic acid were derivatized with o-phenylenediamine to improve mass spectrometric sensitivity. Quality control (QC) samples were prepared by pooling aliquots of study samples and injected after every four study samples to monitor instrument performance throughout these analyses.

The analysis was performed on a Shimadzu 20AD HPLC system and a SIL-20AC autosampler coupled to 4000Qtrap mass spectrometer (AB Sciex) operated in positive multiple reaction monitoring (MRM) mode. Data processing was conducted with Analyst 1.6.3 (Applied Biosystems). The relative quantification data for all analytes were presented as the peak ratio of each analyte to the internal standard. The Metabolomics Facility of Washington University School of Medicine performed the analysis.

### Statistical analyses

Normality was tested using the Shapiro-Wilk test. For normally distributed data, statistical significance was assessed using either Student’s t-test or one-way analysis of variance (ANOVA) with post-hoc Dunn’s test or Student-Newman-Keul’s to compare with the control group. For non-normally distributed data, the Mann-Whitney U-test or the Kruskal-Wallis test were used. All statistical analyses were performed using SigmaPlot 12.5 (Systat, San Jose, CA, USA) or Prism (GraphPad, Boston, MA, USA). In all cases, p<0.05 was considered significant.

## DATA AVAILABILITY

All data are included within this manuscript. Raw data files are available upon request.

## ACKNOWLEDGEMENTS

We thank the Division of Laboratory Animal Medicine (DLAM) at UAMS for expert animal care and Jennifer D. James, BS, HT/ASCP, HTL, QIHC in the Experimental Pathology Core Laboratory for assistance with immunohistochemistry.

## ABBREVIATIONS

ALF: acute liver failure
ALT: alanine aminotransferase
APAP: acetaminophen
CKO: combined ALT2/MPC knockout
GSH: reduced glutathione
GSSG: oxidized glutathione
JNK: c-Jun N-terminal kinase
KO: knockout
LDH: lactate dehydrogenase
MPC: mitochondrial pyruvate carrier
NAPQI: N-acetyl-p-benzoquinone imine
P450: cytochrome P450
PDH: pyruvate dehydrogenase
PDK4: pyruvate dehydrogenase kinase 4
TCA: tricarboxylic acid

## Supplemental Figures

**Suppl. Fig. 1.**
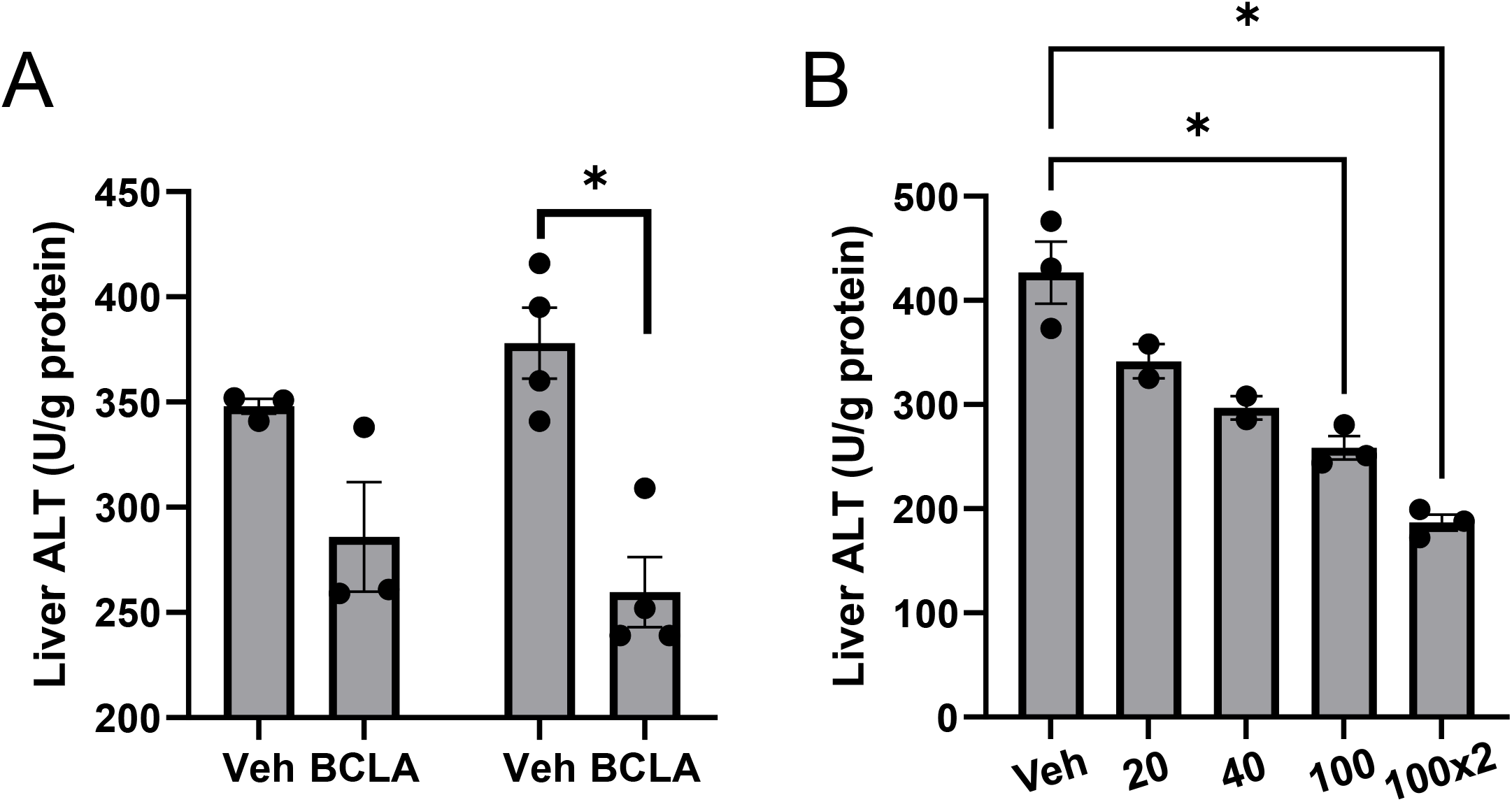
Effect of BCLA on liver ALT activity *in vivo*. (A) WT mice were treated with 20 mg/kg BCLA or vehicle control and total ALT activity was measured in liver tissue lysates 4 (left) and 24 (right) hours later. (B) WT mice were treated with the indicated doses (20 – 100 mg/kg) of BCLA for 24 hours and total ALT activity was measured in liver tissue lysates. Data are expressed as mean±SE. *p<0.05.

**Suppl. Fig. 2.**
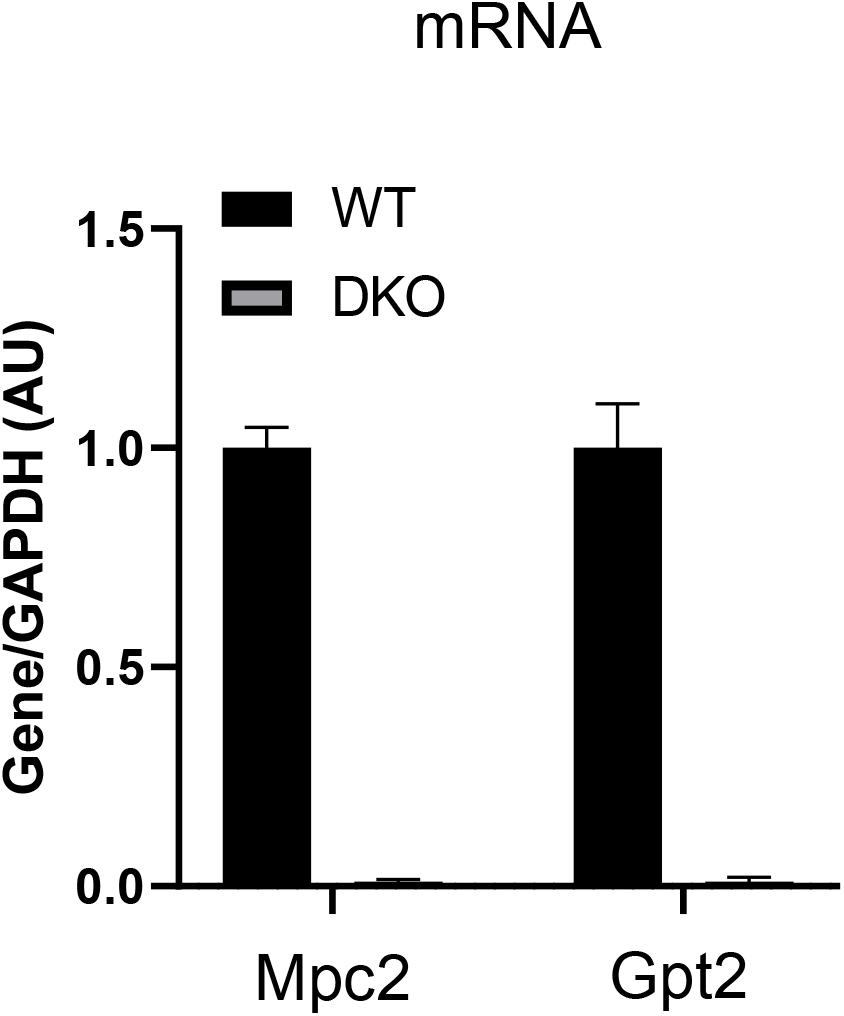
Expression of MPC2 and GPT2 (ALT2) in liver tissue from WT and DKO mice. mRNA for MPC2 and ALT2 was measured by qPCR in RNA isolated from liver tissue and converted to cDNA. Data are expressed as mean±SE. *p<0.05.

**Suppl. Fig. 3.**
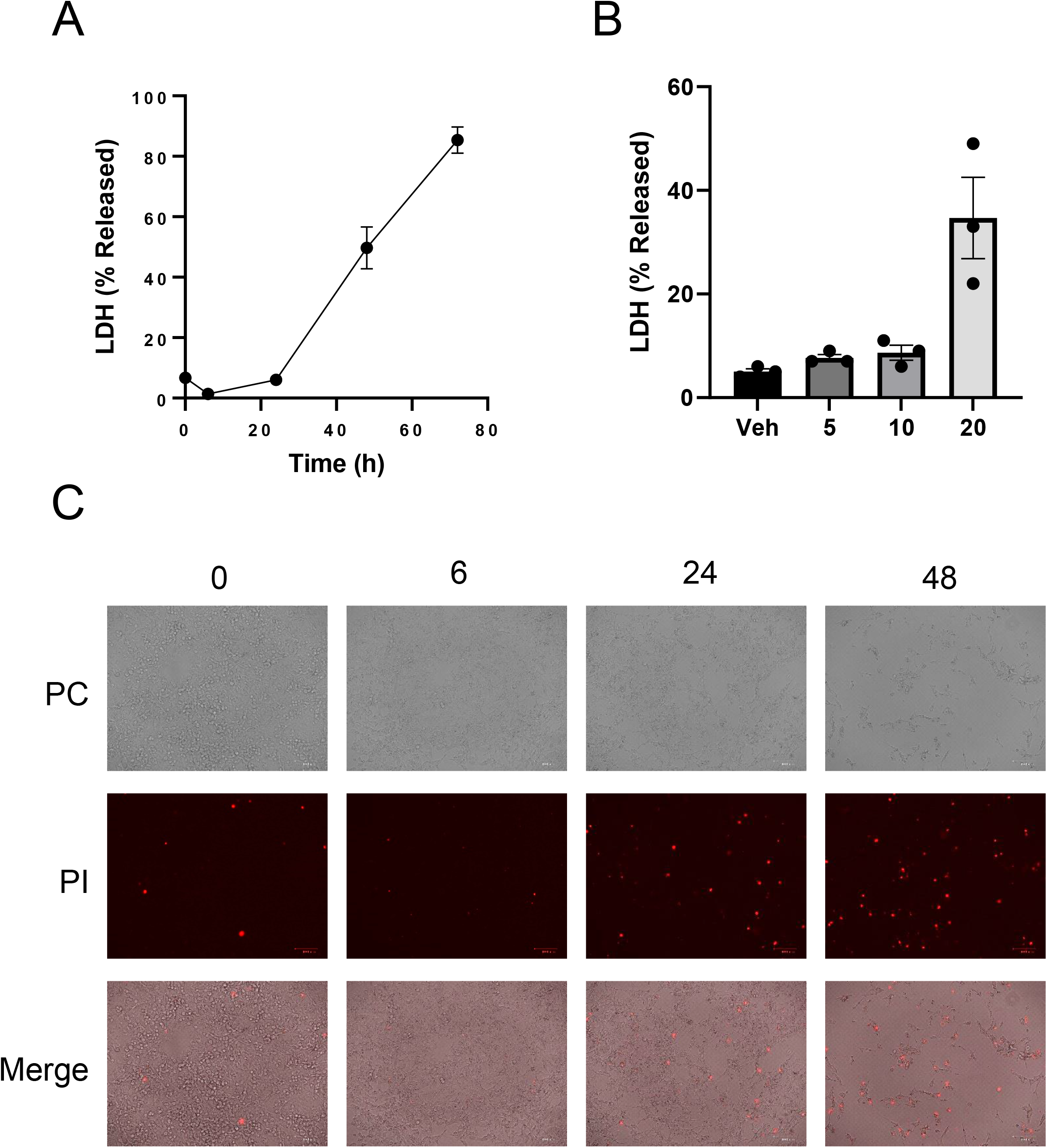
Characterization of APAP toxicity in HepG2 cells overexpressing CYP2E1. (A) Cells were treated with the 20 mM APAP for the indicated time points and LDH release into medium was measured. (B) Cells were treated with the indicated concentrations of APAP for 48 hours and LDH release was measured. (C) Cells were treated with 20 mM APAP for the indicated time points and stained with propidium iodide (PI). Data are expressed as mean±SE for 3 independent experiments run on different days. *p<0.05 vs. Veh control.

**Suppl. Fig. 4.**
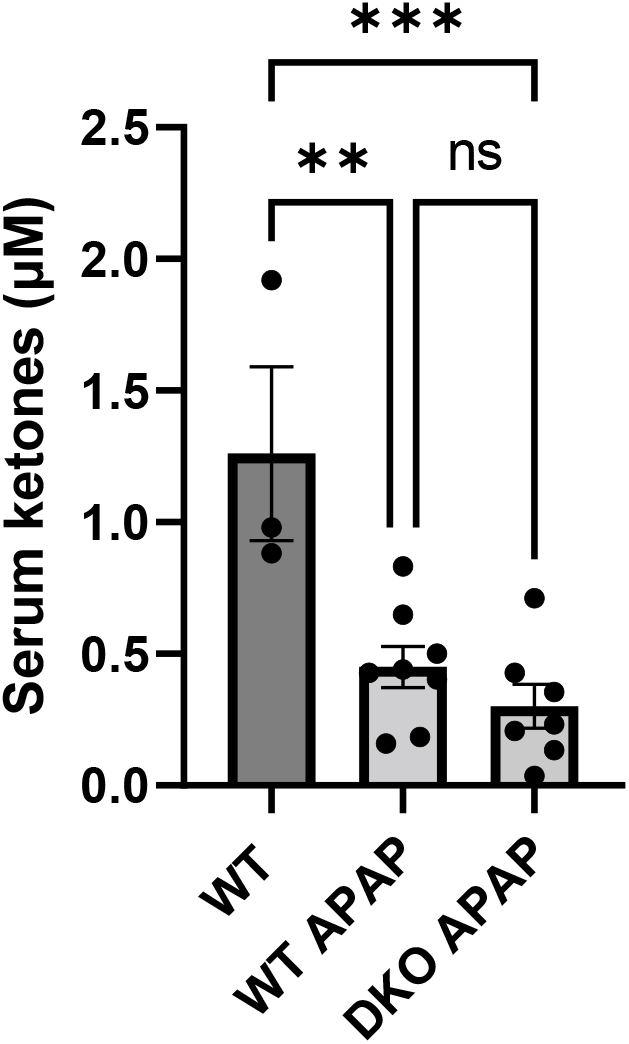
Serum ketones after APAP overdose. β-hydroxybutyrate was measured in serum from vehicle-treated WT mice and from WT and DKO mice after APAP overdose. Data are expressed as mean±SE for 3 independent experiments run on different days. *p<0.05. **p<0.01. ***p<0.001.

